# Global effects of identity and aging on the human sperm methylome

**DOI:** 10.1101/2023.03.21.533698

**Authors:** Guilherme de Sena Brandine, Kenneth I Aston, Timothy G Jenkins, Andrew D Smith

## Abstract

As the average age of fatherhood increases worldwide, so too does the need for understanding effects of aging in male germline cells. Molecular change, including epigenomic alterations, may impact offspring. Age-associated change to DNA cytosine methylation in the cytosine-guanine (CpG) context is a hallmark of aging tissues, including sperm. Prior studies have led to accurate models that predict a man’s age based on specific methylation features in the DNA of sperm, but the relationship between aging and global DNA methylation in sperm remains opaque. Further clarification requires a more complete survey of the methylome with assessment of variability within and between individuals.

We collected sperm methylome data in a longitudinal study of ten healthy fertile men. We used whole-genome bisulfite sequencing of samples collected 10 to 18 years apart from each donor. We found that, overall, variability between donors far exceeds age-associated variation. After controlling for donor identity, we see significant age-dependent genome-wide change to the methylome. Notably, trends of change with age depend on genomic location or annotation, with contrasting signatures that correlate with gene density and proximity to centromeres and promoter regions. These molecular signatures reflect a stable process that begins in early adulthood, progressing steadily through most of the lifespan, and warrants consideration in any future study of the aging sperm epigenome.

## 1 Introduction

Molecular alterations in human germ cell DNA may affect fertility and health risks for the offspring. Several diseases are known to correlate with paternal age at conception. Examples include autism [1], schizophrenia [2], bipolar disorder [3], Huntington’s disease [4] and childhood leukemia [5]. The rates of spontaneous mutation and epigenetic alteration in male germ cells increase with age [6] and are plausible causes for age-associated decrease in fertility and increased risk of disease in progeny. Specific epigenetic features of sperm are strong predictors of viability when embryos are grown in vitro [7], more so than common parameters used in semen analysis, such as volume and motility [8]. Evidence mounts for an epigenomic contribution in connecting paternal age with offspring viability and health.

DNA cytosine methylation refers to the addition of a methyl group on the pyrimidine ring of cytosines. Mammalian DNA methylation occurs predominantly in the cytosine-guanine (CpG) context. Methylation relates to cell phenotype, acting as part of the concerted epigenome to regulate gene expression. The methylation level at a CpG site refers to the fraction of molecules in the cell population with the methyl mark at that site. Methylation is known to have a strong association with chronological age. Using the levels at a small number of CpG sites in almost any tissue, the donor’s age can be accurately calculated, and, surprisingly, the same formula can be used in almost all human cell types with little change in accuracy [9]. Sperm is an exception, suggesting that aging has a different effect on its epigenome. Nonetheless, predictive models trained solely on sperm data can also predict age with similar accuracy to models trained in somatic cells [10]. Various changes to the sperm methylation are known to correlate with both age and fertility in mammals. In mouse, an age-dependent increase in methylation of ribosomal DNA is seen both in sperm [11] and liver cells [12]. Likewise, loss of CpG methylation in coding regions occurs in the sperm of older mice, with similar methylation patterns observed in brain of the offspring [13]. These observations suggest possible age-associated changes to the mammalian sperm chromatin with implications for fertility and offspring phenotype.

To date, the largest population-scale studies of DNA methylation have used array-based technology, a cost-effective and highly reproducible means of studying DNA methylation, but which includes only a fixed subset of CpG sites. Insights from array-based data have been essential in predictive modeling of the epigenome. In contrast, whole-genome bisulfite sequencing (WGBS) provides the richest data to study DNA methylation. WGBS has been more expensive than arrays, but it interrogates the genome at much higher spatial resolution [14, 15]. WGBS has dramatically refined our view of how DNA methylation is organized and how it changes through development, disease progression and aging. Notably, in a homogeneous healthy somatic cell population, the profile of methylation levels along the genome – the methylome – takes the form of mostly high levels punctuated by defined valleys of low methylation. These hypomethylated regions (HMRs) mark promoters and enhancers in a cell type-specific manner, and have a close relation with accessibility [16, 17]. WGBS illuminated a methylome feature prominent in cancers: global hypomethylation of the genome [18]. Early WGBS studies revealed this to be organized in large intervals called partially methylated domains (PMDs), due to their typical intermediate methylation levels. PMDs are associated with regions of heterochromatin, lamina attachment and late replication [19, 20, 21]. We conjectured that WGBS may similarly extend our understanding of the aging sperm methylome, complementing knowledge established in array-based studies.

Sperm cells have unique epigenetic features not observed in somatic cells. Their chromatin is organized by protamines that significantly compact DNA in the sperm head [22]. The sperm methylome harbors additional HMRs at retrotransposons [23]. In many species of mammal, the HMRs at promoters are significantly wider in sperm than in somatic cells [24]. So far, however, limitations in the available human reference genome have restricted our analysis, most notably excluding centromeres. The telomere-to-telomere human assembly provides new opportunities to study the methylome more broadly [25]. Using this assembly, early methylation analyses of human cell lines have revealed large-scale hypomethylation co-localizing with centromeric protein A [26]. The centromeric epigenome may be relevant to aging. Changes to centromere organization have been linked to cell senescence, including an unraveling of satellite heterochromatin [27] and a decrease of centromere protein abundance [28]. Much less is known about how, or if, the centromeric epigenome changes with age in sperm.

More generally, little is known about variation of global features of the sperm methylome within and between individuals. The successes in predictive modeling have, by necessity, emphasized precise features defined on specific sites or small genomic regions. Considerable data normalization is necessary for cross-sample microarray comparisons and this can reduce the apparent variation in these data. The best predictive models of age or clinical outcome implicitly select features with low inter-individual variability. To the extent these models are accurate, the features they rely on must exhibit low variation between individuals. However, we cannot properly study methylome variation – within or between individuals – unless we have repeated measures. In a context where aging is already known to be important, this requires a longitudinal design.

The present study is motivated by two broad questions. First, can high-resolution data reveal additional connections between the sperm methylome and age? Second, are there features of the sperm methylome that vary more between individuals than within? We answer both questions in the affirmative. We show that many global methylome features vary substantially between individuals, regardless of age, while remaining stable long-term within the individual. In other words, the methylome of two sperm samples from the same donor, even taken many years apart, are likely more similar to each other than to an age-matched sample from a different donor. We also identify several methylome features that change consistently with age, across donors. These features are often only appreciable once the effect of individual has been removed, and the total fraction of the sperm methylome impacted by aging is remarkably high.

## 2 Results

### 2.1 High-resolution methylomes in a longitudinal design

We produced 20 WGBS data sets from 10 individuals at 2 time points, separated by 10 to 18 years (Figure 1A). Donors were healthy men, all of whom had healthy children between the collection of consecutive samples. Younger sperm samples were identically preserved in liquid nitrogen with a previously established assay [29] used for *in vitro* fertilization [30].

**Figure 1:**
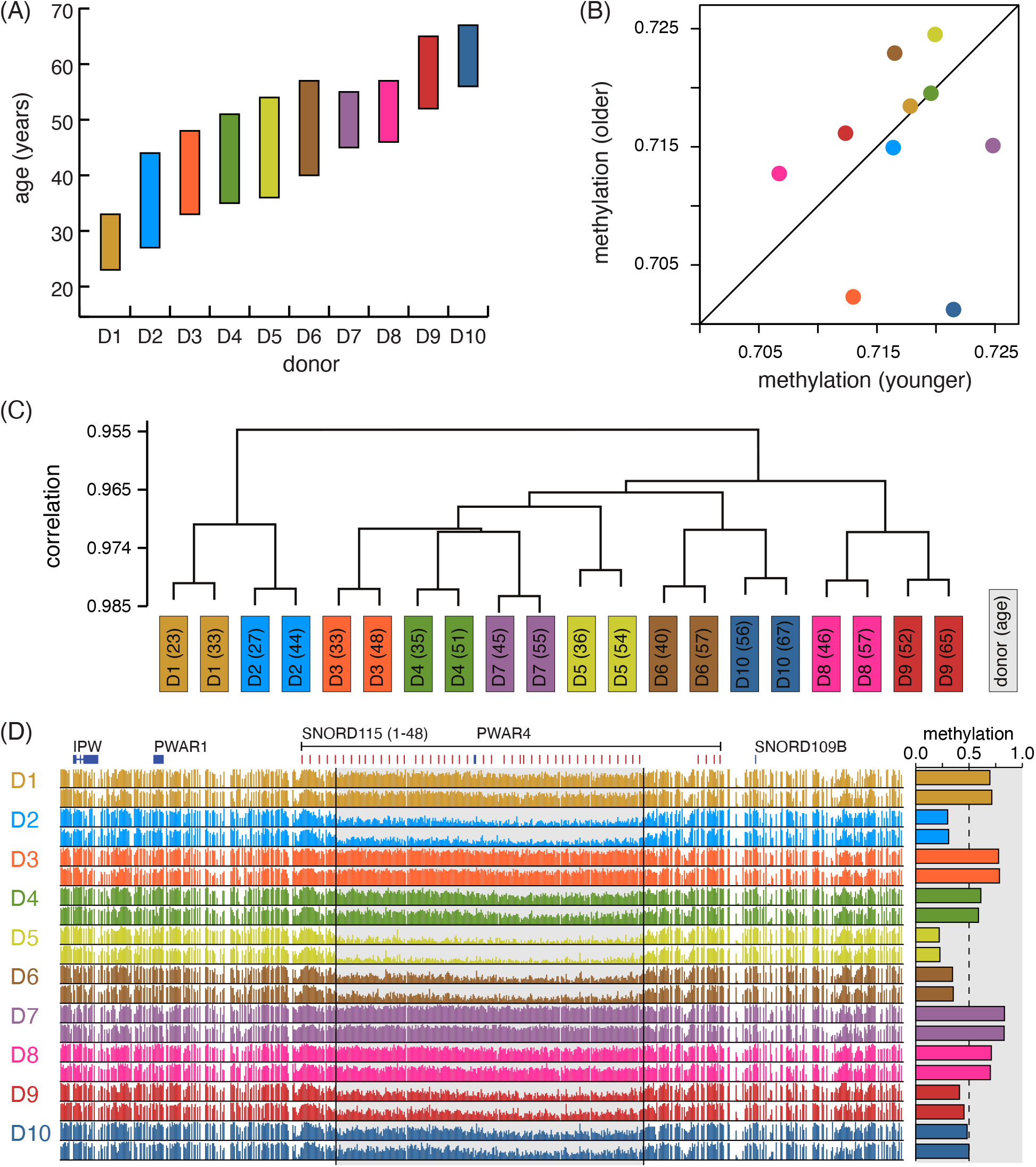
Donor identity dominates global variation in human sperm DNA methylation. (A) Study design includes 10 donors with two sperm samples collected at least 10 years apart from each donor. (B) Genome-wide methylation levels for both samples (“younger” vs. “older”) from each donor. (C) Hierarchical clustering of methylation profiles at 10 kb resolution. (D) DNA methylation through the snoRD115 locus plotted for each sample (younger sample above older); (right) weighted mean methylation through the interval covering snoRD115-5 to snoRD115-44. (All data in this figure are colored as indicated in panel A).

Bisulfite-converted libraries were constructed from sperm DNA and sequenced with Illumina instruments to produce short paired-end reads of 2 × 100 bases. Libraries were sequenced to attain at least 10x coverage of CpGs genome-wide when combining both strands. To our knowledge, this is the first set of WGBS data that covers a comprehensive age range (more than 40 years) of the adult male lifespan.

Sequenced WGBS reads were mapped to the human telomere-to-telomere genome (CHM13 v2.0) [31, 25]. A methylation level was estimated at every cytosine in the genome as the fraction C/(C + T) of read cytosines (Cs) and thymines (Ts) mapping over a reference cytosine [32]. High data quality was confirmed by multiple metrics (Table S1). Reads from all 20 samples had an approximately 80% mapping rate to the human reference genome, with a negligible number of reads mapped to mitochondrial DNA. Bisulfite conversion estimated using cytosines outside the CpG context was above 99% for all samples, which also indicates non-CpG methylation is negligible. There was also no correlation between bisulfite conversion rate and global methylation levels (*p* = 0.3). Methylation levels at CpG sites matched the expected fraction of methylated CpGs reported in prior studies at approximately 70% [10], with a tight distribution (see below) of global methylation level across all 20 samples. These data uniformly interrogate the fraction of the human genome that is mappable by paired-end reads of size 2 × 100 bases.

The global (weighted) average CpG methylation level for all samples varies between 69.2% and 72.0% (Methods). Between different collection times, for the same donor, global methylation level changes range from -0.6% to 1.3%. We found no directional change within donors (Figure 1B, *p* = 0.39, paired *t*-test) or correlation between donor identity and methylation level (*p* = 0.8). Similarly, we did not observe directional change in methylation in annotated CpG islands. We also did not detect a correlation between age and frequency of CpG*→* TpG mutations, within our estimates, although such a correlation likely exists. This indicates that any findings of methylation loss with age would not be an artifact of accumulated C *→* T mutation.

We compared these 20 methylomes on a finer scale by partitioning each in bins of 10 kb, which captures variation in methylation along the genome. We calculated the weighted average methylation level in each bin (Methods). Hierarchical clustering of these profiles shows that methylomes from the same donor exhibit much greater similarity than any pairs from distinct donors (Figure 1C). This implies that, at least for the 10 kb resolution, donor identity is the single strongest factor accounting for variation in the methylomes.

One striking example of DNA methylation determined by donor identity is the snoRD-115 locus, an array of small nucleolar RNAs on chromosome 15 covering roughly 100 kb. This locus is contained in the chromosome region 15q11-13, which overlaps an imprinted gene cluster whose alleles are expressed by parent-of-origin rather than genotype [33]. The deletion of a 2 Mb locus in this cluster from the paternal or maternal allele cause Prader-Willi or Angelman Syndrome, respectively [34]. Aberrant DNA methylation of various genes often co-occur with these deletions, and targeted methylation measurements of these loci serve as a highly accurate diagnostic test for both diseases [35]. It is hypothesized that repression of the SNORD115 gene coupled with post-transcriptional silencing of the serotonin receptors may be responsible for some behavioral features of the syndrome in humans [36]. In our data, the methylation profile of the snoRD-115 locus varies strikingly between donors but remains stable with age within each donor (Figure 1D). The methylation level through roughly 80 kb of this locus (and covering 39/48 snoRD-115 genes) varies from highly methylated (e.g., donor D1) to very low methylation (e.g., donor D2). This example highlights the extent of biological variation between the sperm methylomes of healthy fertile men, and that this can be independent of age.

### 2.2 Centromere methylation accumulates with age

Prior studies of the aging sperm methylome have found relationships focused precisely on individual sites [10]. As a first step to identify parts of the methylome that may exhibit an age effect in our data, we applied a mixed model to relate age and methylation levels summarized in 100 kb bins along each chromosome (Methods). The resulting profiles of conditional *R*^2^ values indicate the relative fit to the data in different parts of the genome (Figure 2A). The most striking feature of these profiles is that they exhibit clear peaks through centromeres, the positions of which are also indicated in Figure 2A. Beyond the positive correlation between methylation levels and age, our data support previous observations of centromere methylation in human sperm: across donors and ages, centromeres are substantially hypomethylated relative to the rest of the genome [11].

**Figure 2:**
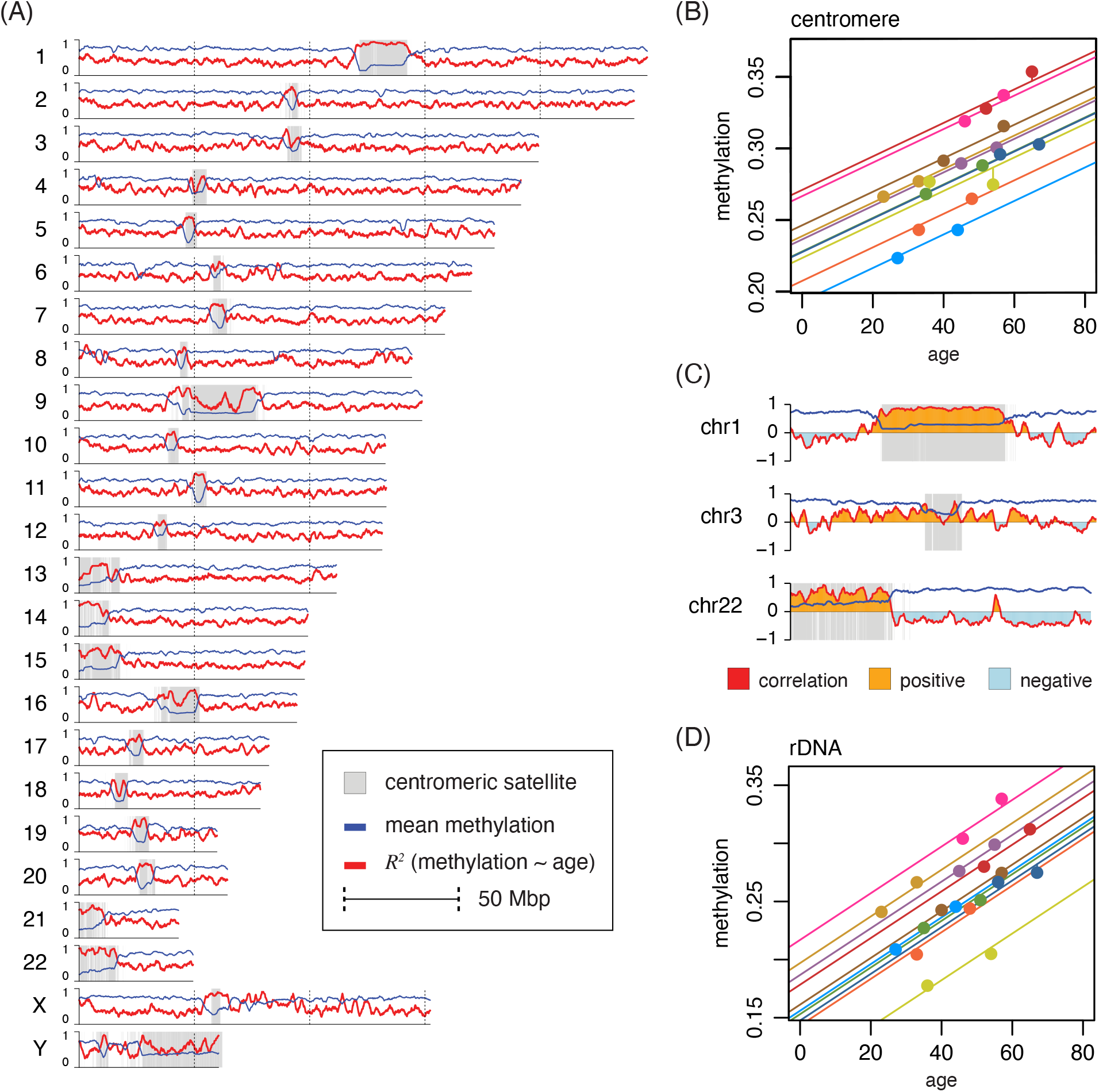
DNA methylation increases with age through centromeres. (A) Methylation and conditional coefficient of determination (*R*^2^) for methylation *∼* age, evaluated in 100 kb bins, along each chromosome. (B) Rainbow plot of methylation as a function of age through centromeric satellites (centromeric transition regions excluded). (C) Correlation of age and methylation plotted through 50 kb of chromosomes 1, 3 and 22, indicating both positive and negative correlation relative to the centromere position. (D) Rainbow plot of methylation as a function of age for ribosomal DNA annotations. (Donors colored as in Figure 1.)

We then directly asked if this visually apparent trend through centromeres, defined via satellite sequences (Methods), is significant. We found a correlation of 0.72 between age and methylation level through centromeres (*p* = 0.0003). However, given the strong effect of donor shown in Figure 1C, we decided to explicitly account for donor identity in our models. We fit a linear mixed model to the data using donor identity as a random effect (Methods) and tested whether this results in a significantly better fit. Our analysis confirm that the mixed model with donor as random effect is appropriate for these data (*p* = 3.2 × 10^−5^). We see a conditional correlation of 0.95 (*p* = 1.5× 10^−4^) for average centromere methylation and age at time of sample collection. We depict this relationship by drawing a separate line for each donor, with all donors sharing a common slope (Figure 2B; Table S2). Lines are colored according to the scheme of Figure 1A, and we refer to this type of plot as a “rainbow plot.” In Figure 2B, all donors except one (D5) have an increase in centromere methylation with age, and the increase with age is very consistent after accounting for each donor’s unique baseline methylation level.

Human centromeres are notably diverse in sequence composition and chromosomal position, thus we do not expect them to behave uniformly. Applying the mixed model approach to individual chromosomes, three did not reach statistical significance at a level of 0.05 (chromosomes 3, 4 and 12). The consistently high conditional *R*^2^ value through the centromere of chromosome 1, seen in Figure 2A, reflects a positive correlation that also drops just outside the annotated centromere (Figure 2C). We can also see that the centromere of chromosome 3 lacks this feature. Although chromosomes 1 and 3 are both metacentric, their centromeres have different sizes: 21 vs. 6 Mb, covering 8.5% vs. 3.0% of the respective chromosome. The centromere of chromosome 3 includes a large human satellite 1A repeat, flanked by active and inactive higher-order repeat (HOR) elements. The centromere of chromosome 1 includes a massive contiguous human satellite 2 repeat, with an uninterrupted active HOR element, together making up almost the entire centromere. Although analyses for families of satellite repeats reveal some trends, in general chromosome identity and centromere composition are too closely coupled to make general claims from our data about which of these two factors may be more important in driving overall trends.

An exception, however, is the acrocentric autosomes (13-15, 21, 22), which all show a strong trend towards increased sperm methylation with age. In human, all acrocentric centromeres contain the ribosomal RNA genes (rDNA), and physically localize to the nucleolus. The positive correlation through the centromere in chromosome 22 is illustrated in Figure 2C, and falls abruptly just after the centromere. We analyzed methylation of rDNA genes, copies of which reside in each acrocentric centromere, but only in these centromeres. We found a correlation of 0.99 after correcting for donor identity (Figure 2D), which agrees with previous similar observations in the human sperm methylome [11]. Although the gain of methylation in acrocentric centromeres with age in human sperm remains strong outside the rDNA, the strongest trend we observe is within those elements.

### 2.3 Replicating timing implicated in global age effects

By visual inspection of the profile in Figure 2A, we noticed that the age vs. methylation relationship drops through several gene-dense loci flanked by gene deserts. We hypothesized that the correlation shows a relative elevation through regions identified as PMDs in somatic-derived cells. To test this hypothesis, we used seven different public methylomes known to contain PMDs (Methods). For each of these methylomes, we identified PMDs (excluding centromeres) and conducted analyses relative to the PMD and non-PMD portions of these methylomes. PMDs in these methylomes covered between 31% and 52% of the genome outside of centromeres, with an average of 41%.

We first asked how the age vs. methylation correlation, shown in Figure 2C, relates to PMD status. For each of the seven PMD sets we computed the distribution of these estimated correlation coefficients (Figure 3A). The PMD and non-PMD portions have strikingly different distributions: the PMD portion appears roughly symmetric, while the non-PMD portion of the genome skews negative, with a mode significantly below zero. The age-associated gain of methylation in PMDs is consistent with our observation in centromeres. Notably, the majority of bins that gain methylation (with *p <* 0.01) lie in PMD regions and, conversely, the majority of bins that lose methylation are in non-PMD portion (Figure 3B). Similar to PMDs, centromeres are associated with late replication in human cells [37].

**Figure 3:**
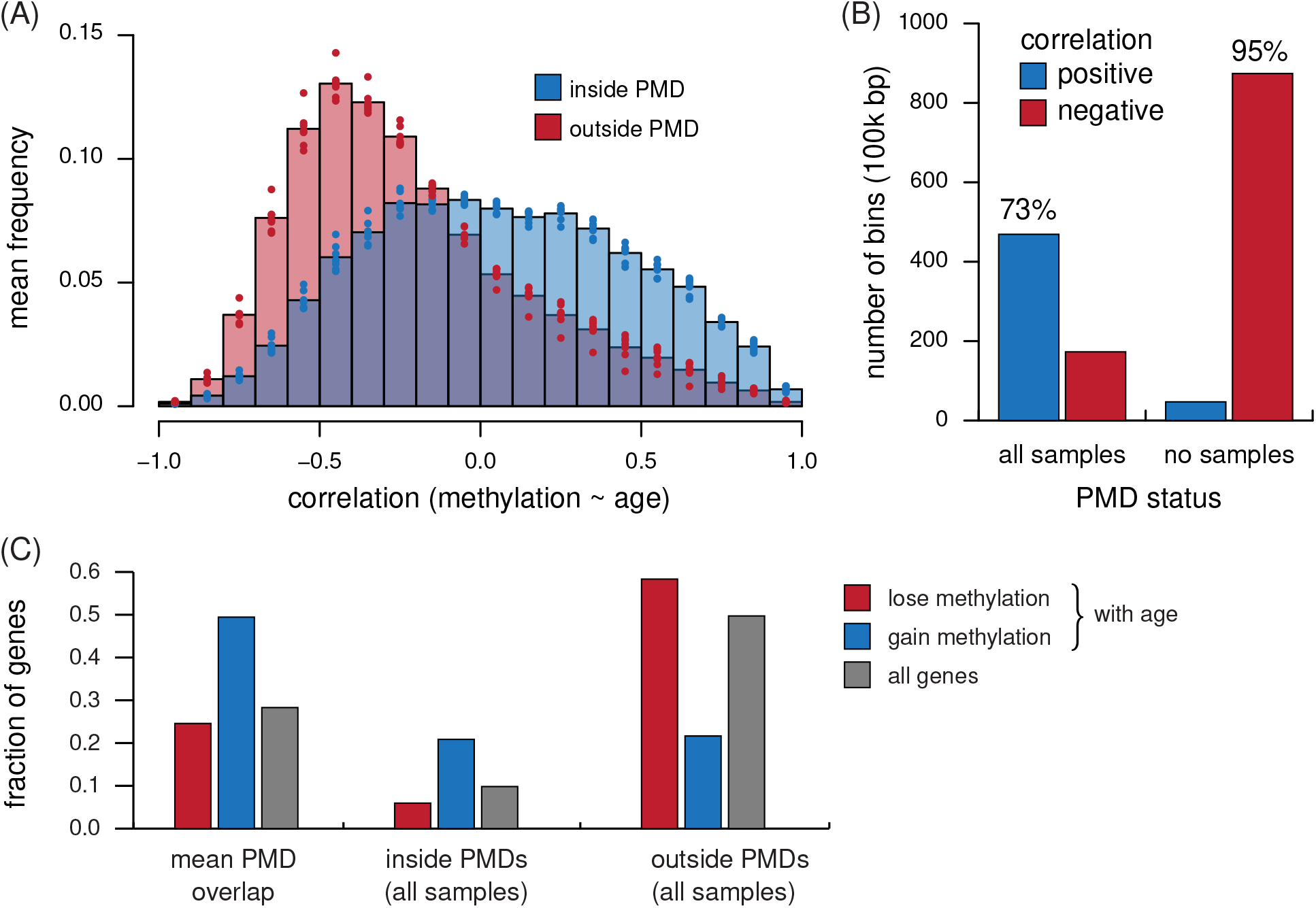
Age effects in estimated early- and late-replicating regions of the genome. (A) Distribution of age vs. methylation correlation in 100 kb bins inside (blue) and outside (red) of PMDs. Each bar’s height is the mean from 7 PMD-containing methylomes, indicated separately as dots. (B) Number of bins with significant positive (blue) and negative (red) correlations between age and methylation in two groups: bins that overlap a PMD in all (left) vs. none (right) of the methylomes tested. (C) Fractions of genes overlapping a PMD in all or none of the methylomes, separated by genes losing methylation with age in human sperm (red), genes gaining (blue), and the expectation based on all genes (gray). Bar groups: the mean gene overlap with PMDs across all samples (left), genes overlapping regions that are always PMDs (middle) and genes overlapping regions that are never PMDs (right)

The contrasting changes in methylation inside and outside of PMDs might explain previous observations of age-related DNA methylation changes in sperm. Among a published set of 428 genes with methylation level in sperm that can predict age, 172 gained and 256 lost methylation [38] (Table S3). We asked if the stratified correlations inside and outside of PMDs can explain the previous findings of gain or loss with age in these genes. We determined which genes overlap PMDs in each of the PMD-containing methylomes. Among genes reported as gaining methylation with age, on average half overlap a PMD in any given methylome and roughly 20% overlap a PMD in all tested methylomes (Figure 3C). Conversely, most genes that lose methylation with age never overlap a PMD in any tested methylomes. This contrast is also seen when counting nucleotides to control for differences in gene sizes (data not shown). Thus, on the scale of full gene loci, our data connect directional age-associated methylation changes in sperm with the shared structural, replication timing and epigenomic features of PMDs.

### 2.4 Hypomethylated regions disappear with age

The changes we see in centromeres and PMDs, which are connected with replicating timing, are measured on the scale of mega-bases. At higher resolution, methylome features are usually associated with gene regulation. In particular, across most mammalian cell types hypomethylated regions, evident as valleys in a background of high methylation, typically mark active or accessible regulatory regions like promoters and enhancers. HMRs are contiguous intervals on the order of hundreds to a few thousand base pairs. Typical human sperm methylomes have about 70k of these HMRs and average 1.5 kb in size [23].

We identified HMRs in each of the 20 sperm methylomes. We excluded centromeres as the extreme hypomethylation at mega-base scale would confound identifying smaller intervals (Methods). This also ensures that our observations are independent of the age-associated changes outlined above. Across all samples, the number of identified HMRs varies between 55k and 74k, with a mean size in the range of 1,723 to 1,848 bases. We used a similar mixed model strategy as outlined above to determine whether the sets of HMRs change with age. We observe a striking global decrease with age in the number of HMRs (*p* = 1.1 × 10^−5^). This is a loss of 88.7 HMRs per year in the human sperm methylome (Figure 4A; Table S4). Globally we do not observe a significant change to the mean size of regions (*p* = 0.0255). These results indicate that aging has a global effect on features that are likely to impact gene regulation at some stage in sperm development.

**Figure 4:**
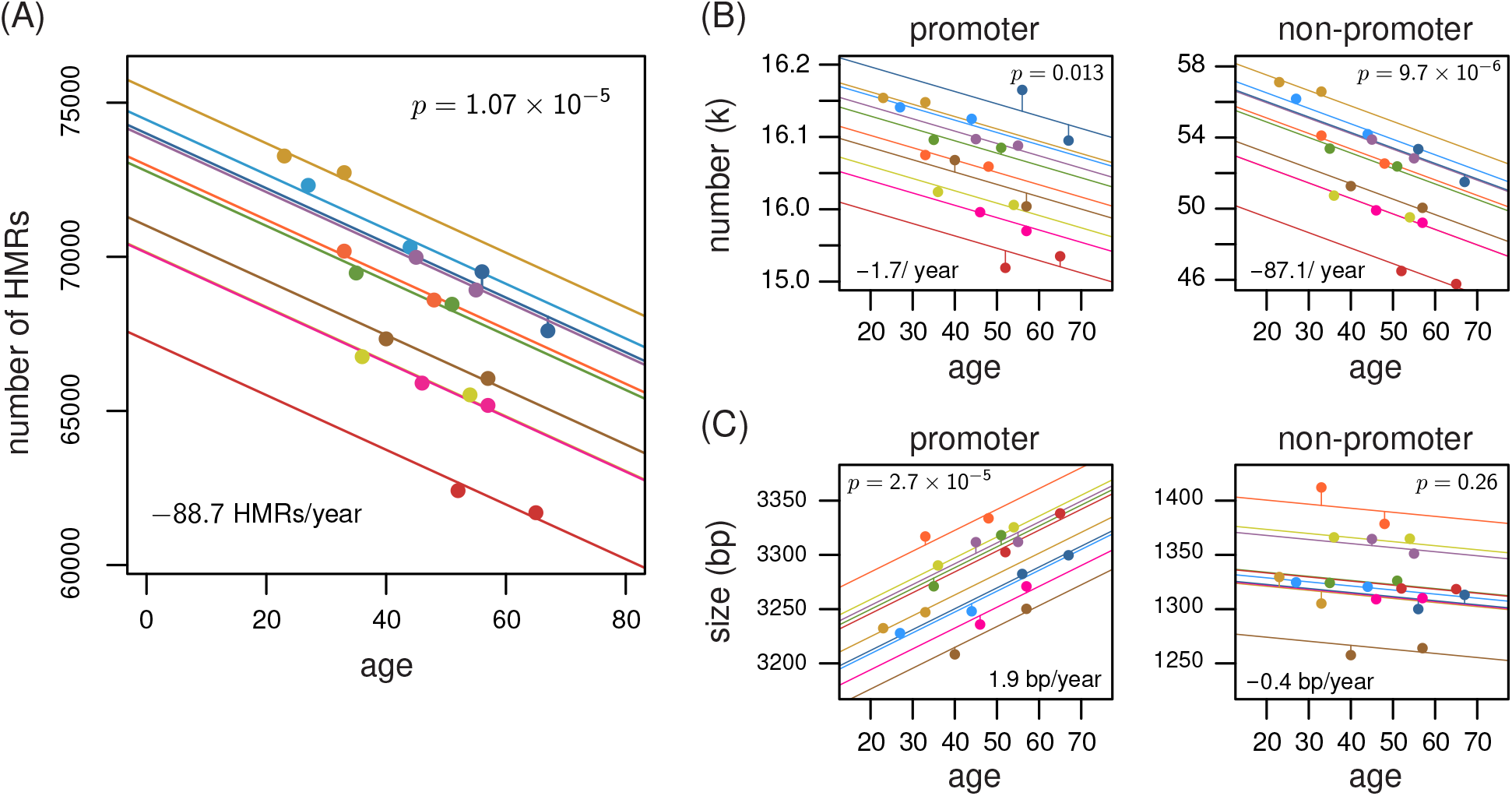
Hypomethylated regions disappear with age. (A) Rainbow plot showing the genome-wide loss of hypomethylated regions with age. (B) Rainbow plot decomposing the trend indicated in panel A according to whether the regions overlap annotated promoters. (C) Rainbow plot showing size of hypomethylated regions as a function of age, for each individual, stratified by whether the regions overlap annotated promoters. (Donors are colored as in Figure 1.)

HMRs in sperm of many species of mammal exhibit unique properties at promoters. Especially for primates, the size of HMRs spanning transcription start sites are substantially wider in sperm than in somatic cells, and within sperm they are wider at promoters than distal regulatory regions or retrotransposons [24]. We repeated the mixed model analysis but with HMRs separated based on whether they overlap a transcription start site (Methods). The proportion of HMRs that overlap transcription starts varies between 23.8% to 27.6%, but the number of such HMRs does not decrease significantly with age (*p* = 0.013; Figure 4B). The average of 25.6k unique promoters covered by an HMR spans a range of only about 1% of all promoters. The majority of HMRs are not at annotated promoters (74-78%), and these reduce significantly in number with age, at a rate of 87 HMRs per year. Therefore, the global decrease in HMRs with age is driven by HMRs away from promoters.

We examined HMR size as a function of age using the same stratification by promoter status and found that the HMRs at promoters do show a significant change in size with age (Figure 4C). In particular, the average size of these HMRs at promoters grows by roughly 1.9 bp per year. The same does not hold for HMRs away from promoters. We can thus conclude that the HMRs at promoters grow wider with age, and HMRs away from promoters disappear with age. In both cases the mixed model indicates a consistent change with age, regardless of the donor’s baseline.

## 3 Discussion

We generated WGBS data from human sperm in a longitudinal design, covering more than 40 years. Genome-wide DNA methylation analyses revealed substantial variability between donors that remains stable with age within the individual. After accounting for donor identity, we observed age-associated changes to the sperm methylome throughout the genome. Centromeres gain methylation with age. Age-associated changes outside centromeres show strong directionality as a function of replication timing and nuclear organization for the underlying genome, as indicated through correlation with previously identified partially methylated domains. Finally, hypomethylated regions, which tend to mark promoters and enhancers, disappear with age when distal from promoters, but are retained and expand in size when localized to promoters. Most of these findings are obscured by inter-individual variation, but emerge prominently after controlling for donor identity.

Centromere sequences encode information that is essential to coordinating chromosome segregation during cell division. The epigenome has a role in this process, but it is unclear whether centromeric DNA methylation directly impacts this process or is a passenger influenced by other components of the integrated epigenome [39]. Our results indicate that centromere methylation, which is reduced compared to the rest of the genome, increases with age in sperm. This phenomenon is detectable even without controlling for donor effect, so the molecular determinants of methylation through centromeres likely degrade with age in a way that differs from most of the genome. Centromeres are notoriously difficult to analyze using shortread sequencing. The telomere-to-telomere reference is a dramatic step forward, but it does not alleviate all technical issues. We analyzed centromeres using aggregate measures. More precise claims would require different approaches – among them, use of long reads [26]. It seems reasonable that within the large human centromeres, higher resolution analyses may reveal detailed patterns, possibly associated with specific sequence features.

We see trends in contrasting directions through portions of the genome labeled PMD and non-PMD: the PMD portion exhibits significantly more age-associated increase to DNA methylation in sperm, while the opposite is seen in the non-PMD portion. These divergent trends agree with current understanding of PMD formation in senescent cells. PMDs represent on average 40% of the genomes in which they are observed [40]. The PMDs in different tissues are very similar, and there seems to be at least a common intersection that comprises 25% of the human genome [41]. This constitutive set includes the largest domains, where DNA replicates latest [42]. PMD formation has been previously linked with aging via the observation of early PMD signatures in centenarian T-cells [43]. PMDs, as originally characterized, have not been seen in sperm. We conjecture that late-replicating regions, which correspond to constitutive PMDs, harbor hotspots for age-associated changes to the sperm methylome. Our findings in mature sperm likely reflect events from all progenitor stages that propagate to the mature sperm. Late-replicating DNA is more prone to mutation [44]. Our results suggest these regions also see an accumulation of error in methylome maintenance. The constitutive PMDs and the centromeres replicate very late during mitosis. Together these observations are consistent with a model in which diffuse accumulation of epigenomic noise in male germ cells depends on replication timing or large-scale features of chromatin structure.

We found that a man loses on average 88 HMRs per year in his sperm, likely with high variability between individual spermatozoa, and this rate seems very stable through the lifespan. DNA methylation correlates strongly with gene expression. In general, genes with methylated promoters are silenced and methylated enhancers or cis-regulatory modules are rendered inactive. Unmethylated promoters tend to be accessible, so their activity depends on the presence of transcriptional regulators. One way epigenomic change at the scale of HMRs might relate to aging is by disruption of transcription regulation for genes involved in specific pathways. Even if such alterations are distributed randomly in the genome, and not targeting specific genes or pathways, they could impair function of an increasing number of genes in an increasing number of cells over time. Importantly, age-associated methylation changes may be reversed post-fertilization. The mammalian methylome is reprogrammed during early embryogenesis [45]. The embryonic methylome is erased and DNA is subsequently remethylated, but regions like imprinted loci, ribosomal DNA and some retrotransposons escape these processes [45, 12]. Phenotypic effects in the offspring require that changes occur in loci that escape methylation erasure events.

Age can be accurately predicted using DNA methylation in sperm measured at a small and specific set of sites. While WGBS provides higher spatial resolutions, presently it estimates methylation at each CpG using far fewer molecules, which limits the power of WGBS to detect subtle changes at individual sites. We found broad variability between individuals that overshadows age-related change. This does not conflict with the accuracy of epigenomic aging calculators based on microarrays. The sites selected when training aging calculators might simply be those least affected by individual variation. Our results complement these previous studies and suggest that the predictive accuracy in existing aging calculators may be further improved by directly accounting for relative stability of features across individuals. The prospect of using WGBS as the basis for predicting age of a sample, using only shallow sequencing, depends on some way to determine individual baselines.

Interpretations for the phenotypic consequences of the methylation variation we report are limited: donors in our study were phenotyped as healthy and fertile. We have, however, established a lower bound on the variation in sperm methylomes among healthy fertile men. Inter-individual variation, especially striking examples like the snorD-115 locus, may have little or no clinical relevance. However, the differences at snorD-115 are dramatic. This variability is comparable to those seen in cancer-normal studies in magnitude and scale. To find a phenotype connected to snorD-115 methylation in sperm would not be surprising. Methylation through the snorD-115 locus is remarkably stable within each donor. Among healthy fertile men, not all sperm in a sample are equally viable. From our data we cannot say if the methylation levels through the snorD-115 locus reflect a homogeneous sperm population or a mixture of epigenotypes. In either case, our data point to an equilibrium methylation level. If methylation in sperm through this locus is genetically determined, we expect stability with age. However, we cannot rule out possible effects of lifestyle as cause for any variability we see between donors. The sperm methylome is known to be affected by smoking [46], exercise [47] and exposure to trauma [48], among various other possible covariates.

Our study highlights the nature of individual differences in sperm methylomes and how they are shaped by aging. Individuals vary substantially at certain loci and through certain epigenomic features. Yet, even among features that exhibit striking differences between individuals, we find remarkable stability within individuals. We used repeated measures and explicitly modeled donor identity, allowing continuous signatures of aging to emerge. The resulting picture is a sperm methylome impacted almost genome-wide by aging. Although each man presents a unique sperm methylome as a baseline, our data suggest aging acts on the sperm methylome according to processes that are invariant with time and uniform across individuals.

### Data availability

Source code with data and R code to reproduce results and figures for this manuscript can be found at github.com/smithlabcode/sperm-methylome-aging-wgbs. Bisulfite sequencing libraries were deposited on the Gene Expression Omnibus (GEO) [49], under accession number GSE222340.

Whole genome bisulfite sequencing for partially methylated domains were obtained from the Sequence Read Archive (SRA) [50] with the following accessions: SRX026834 (adipose-derived stem cells), SRX026835 (adipose grown from adipose-derived stem cells), SRX8582072 (Calu1), SRX038781 (foreskin fibroblasts), SRX276179 (GM12878 cell line), SRX026829 (IMR90 cell line) and SRX323156 (placenta).

## Methods

### Library preparation

DNA was extracted and purified for all sperm samples using the Qiagen DNA extraction kit, after which it was sonicated to generate 100-300 bp fragments. The resulting fragments were end-repaired before adapter ligation, then treated with sodium bisulfite using the Zymo EZ DNA Methylation Gold kit. The resulting DNA was then amplified with PCR and sequenced to generate 100 bp paired-end Illumina reads.

### Data processing and methylome quantification

The data consists of twenty paired-end pairs of FASTQ files with reads of length 100 bp on each end. The quality of reads was assessed using Falco version 1.2.1 [51], which was used to test average PHRED quality scores and base content of each set of sequences. Adapters were trimmed using trim-galore version 0.6.7 [52]. The adapter-trimmed reads were mapped to the CHM13 human genome reference version 2.0 using abismal version 3.1.0 [31]. We created two independent sets of mapped reads files in SAM format. The first contained only uniquely mapped reads, and the second contained uniquely mapped reads and a randomly assigned location for reads that mapped ambiguously. The first set was used anytime we showed or referenced a specific part of the genome (e.g., Figure 1D, Figure 2A). The second set was used to summarize weighted average methylation in large regions, including centromeres and rDNA genes, repetitive regions of the genome (see next section). In both cases, reads were mapped by setting the *c* parameter in abismal to 50,000 to maximize the sensitivity of the mapping algorithm. The mapped reads were analyzed using DNMTools version 1.2.1 [32]. Specifically, paired-end mapped read locations from the output SAM file were converted to single-end using the dnmtools format command. The resulting formatted SAM file was sorted using SAMTools version 1.13 [53]. PCR duplicates, defined as identical reads that map to identical locations in the genome, were removed using the dnmtools uniq command. Hypomethylated regions (HMRs) were identified using the dnmtools hmr command. We used the RefSeq gene annotation [54] from NCBI for the CHM13 reference genome version 2.0. The annotation was used to count genes that overlap PMDs and HMRs. Genes were defined as contained in PMDs if they have a non-empty overlap with any PMD interval. HMRs were classified as promoter HMRs if they have a non-empty overlap to the transcription start site of any gene and non-promoter HMRs otherwise.

### Methylation quantification in annotated regions

We used counts of methylated and unmethylated reads in each CpG to create weighted methylation averages, either genome-wide, or when restricted to one or a set of genomic intervals. This includes equally sized bins of sized 10 kb or 100 kb, as well as regions annotated as centromeric satellites, ribosomal DNA, PMDs and genes. In all cases, the input regions to be used for quantification were given as browser-extensible data (BED) files. BED files of equal-sized bins tiling the genome were created using BEDTools [55] with the CHM13 reference genome chromosome sizes as input. Regions defined as PMDs were inferred using the dnmtools pmd command [41]. Genes and centromeres were retrieved from the UCSC Table Browser, based on data provided by the telomere-to-telomere consortium [56]. For our use, the set of of centromeric intervals was defined as the union of all centromere BED files with the exception of centromeric transition regions (Table S5). In other words, centromeres are defined as the union of the following satellite repeats: active, inactive and divergent *α*-Sat higher-order repeats (hor and dhor), monomeric *α*-Sats (mon), classic human satellites 1A, 1B, 2 and 3 (hsat1A, hsat1B, hsat2 and hsat3), beta and gamma satellites (bsat and gsat), and other annotated centromeric satellites (censat). When we excluded centromeric regions, to be conservative we also excluded the ct annotations.

A methylation level for an interval in a BED file is assigned as a weighted average methylation, as previously suggested [57]. Given a genomic region with *M* CpGs, where CpG *i* is covered by *N*_*i*_ reads of which *n*_*i*_ are methylated (0 ≤ *n*_*i*_ ≤ *N*_*i*_), the weighted average methylation *w* is given by

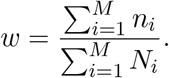

Note that the value of *w* does not depend on where reads map within the region. Through this definition, reads that map ambiguously in the genome but uniquely within a genomic annotation category (e.g., rDNA) can contribute towards estimates of a region’s weighted methylation estimate. We used this property to calculate methylation levels more accurately in highly-repetitive regions, like centromeres and subsets of centromeric satellites. This is analogous to mapping reads to consensus sequences for a family of repeats.

We used unique and ambiguous reads to calculate methylation in centromeres and rDNA. For bins, PMDs, genes and HMRs, we used only uniquely mapped reads. For centromeric regions, unique reads cover 26% of centromere CpGs at 7.6x coverage, whereas unique and ambiguous reads covered at least 47% of CpGs with similar coverage. For rDNA, unique reads covered on average 2% of CpGs, whereas unique and ambiguous reads covered at least 4.92% of CpGs in all samples.

### Comparing statistical models for age-association

Let *y* be a vector of *n* random variables with values estimated using DNA methylation data. Associated with each entry in *y* is an age, which is encoded in the known matrix *X* along with an intercept column. Let *Z* be a known *n* × *m* binary matrix that indicates, for each observation, the corresponding donor. These are related by the following model:

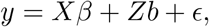

where *β* = (*β*_1_, *β*_0_)^*T*^ gives the unknown fixed effects, with regression coefficient *β*_1_ for the predictor and *β*_0_ gives the grand intercept. The *m* × 1 vector *b* of random effects distributed as 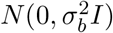. The error *ϵ* is *n* × 1 and distributed as 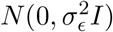 and independent of *b*. This model makes the assumption that individual donors are independent. In our setting, we have *n* = 20 outcomes taken from *m* = 10 individuals, and each of these individuals provided 2 samples, separated by at least 10 years. Thus, within binary matrix *Z*, each row sum is 1 and each column sum is 2. Our null hypothesis is as follows:

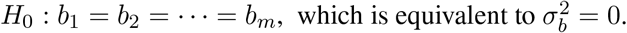

Testing this null amounts to asking whether the data should be modeled using a common intercept for all *m* individuals.

To test *H*_0_, for a particular measurement from the methylome data, we used the exactLRT function from the RLRsim R package [58], which is based on the finite sample distribution of the restricted likelihood ratio test statistic [59]. If we reject *H*_0_, we used the mixed model to test for an additional hypothesis concerning an association between the covariate in *X* and age of the sample, conditional on each donor having a different intercept *b*_*i*_ but shared slope *β*_1_. The null hypothesis for the association with age is

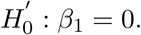

We tested this hypothesis using the lmer function through the lmerTest R package for estimating parameters and hypothesis tests on our fixed effect [60]. To obtain a conditional coefficient of determination (*R*^2^) from a mixed model, as shown in Figure 2A along each chromosome, we used the method of [61], implemented in the MuMIn R package.

### Partially methylated domain datasets

We used partially methylated domains of seven samples, including primary cells and cell lines, as replicates to create a genome partition between early and late replicating regions. We chose the following cell lines and phenotypes (1) adipose-tissue derived stem cells (ADS) (2) ADS-derived cultured adipose (3) The Calu1 squamous cell carcinoma cell line (4) foreskin fibroblasts (5) the GM12878 cell line (6) the IMR90 cell line and (7) placenta. Public SRA accessions for each of these samples are listed in the Data Availability section in the form of Sequence Run Experiment (SRX) IDs. Public datasets were processed using the same software tools and parameters as the sperm samples. [40, 17, 62] (Data Availability)

